# Scaling and Merging Time-Resolved Laue Data with Variational Inference

**DOI:** 10.1101/2024.07.30.605871

**Authors:** Kara A. Zielinski, Cole Dolamore, Harrison K. Wang, Robert W. Henning, Mark A. Wilson, Lois Pollack, Vukica Srajer, Doeke R. Hekstra, Kevin M. Dalton

**Affiliations:** School of Applied and Engineering Physics, Cornell University, Ithaca, NY 148532; Department of Biochemistry and the Redox Biology Center, University of Nebraska, Lincoln, NE 68588; Department of Molecular and Cellular Biology, Harvard University, Cambridge, MA 02138; Graduate Program in Biophysics, Harvard University, Boston, MA 02115; BioCARS, Center for Advanced Radiation Sources, The University of Chicago, Lemont, IL 60439; John A. Paulson School of Engineering and Applied Sciences, Harvard University, Cambridge, MA 02138; Linac Coherent Light Source, SLAC National Accelerator Laboratory, Menlo Park, 94025, CA, USA; Department of Biology, New York University, New York, NY 10003

**Author notes:** (Electronic mail), (Electronic mail), (Electronic mail) (Vukica Srajer, Doeke R. Hekstra, Kevin M. Dalton).

## Abstract

Time-resolved X-ray crystallography (TR-X) at synchrotrons and free electron lasers is a promising technique for recording dynamics of molecules at atomic resolution. While experimental methods for TR-X have proliferated and matured, data analysis is often difficult. Extracting small, time-dependent changes in signal is frequently a bottleneck for practitioners. Recent work demonstrated this challenge can be addressed when merging redundant observations by a statistical technique known as variational inference (VI). However, the variational approach to time-resolved data analysis requires identification of successful hyperparameters in order to optimally extract signal. In this case study, we present a successful application of VI to time-resolved changes in an enzyme, DJ-1, upon mixing with a substrate molecule, methylglyoxal. We present a strategy to extract high signal-to-noise changes in electron density from these data. Furthermore, we conduct an ablation study, in which we systematically remove one hyperparameter at a time to demonstrate the impact of each hyperparameter choice on the success of our model. We expect this case study will serve as a practical example for how others may deploy VI in order to analyze their time-resolved diffraction data.

## I. INTRODUCTION

Time-resolved X-ray crystallography enables the observation of macromolecular dynamics at atomic resolution, capturing essential information to understand reaction mechanisms. The applications of this technique are extremely broad, and within this paradigm, many experimental approaches have been developed at both synchrotrons and X-ray free electron lasers (XFELs). These include pump-probe^1–3^, mix-and-inject serial crystallography (MISC)^4–7^, temperature jump^8^, electric field stimulation^9^, and others, which allow researchers to observe conformational changes of diverse biological targets^10,11^. In this manuscript, we focus on the data analysis of a successful MISC Laue diffraction experiment carried out at the Advanced Photon Source Sector 14 (BioCARS^12,13^). This experiment investigated the reaction of a human enzyme implicated in early-onset Parkinson’s disease, DJ-1, with one of its putative substrates, methylglyoxal. While this is a fascinating biochemical system of considerable medical relevance, here we will focus on the challenges inherent in extracting time-resolved signal from X-ray diffraction experiments rather than the biological conclusions of our study. Based on our experience, we offer practical advice to those conducting time-resolved and comparative crystallography experiments.

The analysis of time-resolved diffraction data is still frequently challenging. The experiments themselves introduce many sources of variability. The goal of any time-resolved experiment is to initiate dynamics as uniformly as possible for all of the protein molecules inside a crystal. Then, assuming sufficient initiation and reaction progress (which can be a challenge for optical pump-probe experiments due to low quantum efficiencies or sample absorbance), small structural changes occur and are captured at predetermined delay times with X-ray pulses. During data collection, the average structure in time across the crystal is recorded. Many time-resolved measurements are coupled to serial crystallography experiments, where datasets comprise up to hundreds of thousands of crystals, introducing additional challenges, such as heterogeneity across microcrystals and the need for accurate scaling and merging of reflections across the dataset. Time-resolved differences are most commonly analyzed with difference electron density (TR-DED) maps. The Fourier coefficients of such a map consist of the difference in structure factor amplitudes between two time points. The phases are usually approximated by those of a high-resolution reference structure of the crystal in the ground state. While DED maps are exquisitely sensitive to changes in the composition of crystals^14^, they are equally sensitive to measurement errors^15^. The success of a time-resolved crystallography experiment may therefore hinge upon the accuracy with which differences in structure factor amplitudes can be measured, which is a noteworthy challenge. The changes in structure factor amplitudes between time points are, in most cases, small relative to the structure factors themselves^14^. This means that data must be measured with unusually high precision in order to accurately infer differences in structure and observe intermediate states. Contamination of the structure factor differences by outliers can mask signal in otherwise well-measured data^15^.

Serial Femtosecond Crystallography (SFX) experiments were pioneered at X-ray Free Electron Lasers^16^, and Serial Synchrotron Crystallography (SSX) experiments are becoming increasingly popular, with both monochromatic^17–20^ and polychromatic X-rays^21,22^. Data collection from quasi-monochromatic sources, however, record only partial Bragg reflections. The most common strategy to handle this partiality, and thus achieve more precise estimates of structure factors, is serial crystallography at extreme statistical redundancy^23^. Our experimental design is different, and exploits a polychromatic synchrotron beamline (BioCARS 14-ID, Advanced Photon Source), which has already been used for previously successful SSX experiments^21,24,25^. The increased reciprocal space information available in polychromatic diffraction patterns, as well as recording of full rather than partial intensities (without crystal rotation), as compared to monochromatic images enabled us to successfully measure time-resolved changes in structure from fewer microcrystals than are typically required in an SFX experiment. Other synchrotron beamlines are starting to broaden their X-ray bandwidth to 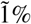 due to these benefits^26–28^. Nonetheless, many of our conclusions will be applicable to time-resolved crystallography experiments, regardless of the X-ray bandwidth or source. The central insight we wish to convey in this manuscript is that recent advances in data analysis can address the measurement errors inherent in crystallography experiments and generate high-quality DED maps. More specifically, we will demonstrate that the statistical method of variational inference^29,30^ is well suited to this challenge.

X-ray diffraction data typically contain many redundant observations of reflections. Merging is the process of averaging the intensities of redundant measurements to estimate a consensus set of intensities which can be used in structure determination or, indeed, Fourier difference map analyses. This task is complicated by the nature of the diffraction experiment, as discussed above. The observed intensity is always modulated by a number of effects which are unavoidable and lead to systematic errors in the measurements. These errors cannot be accounted for strictly by analytical correction factors^31^. Prior to merging, observed intensities are corrected using numerical optimization, a procedure known as scaling^32^. Conventionally, scaling algorithms work by fitting a model of systematic errors in order to minimize the discrepancy between repeated observations of the same reflection. This has been a very successful strategy for conventional, rotation-method data. Every major crystallography software package implements a scaling algorithm. An alternative to sequential scaling and merging was recently proposed which jointly estimates merged structure factors alongside the systematic error model using variational inference^33^. This algorithm is implemented in the Python package, `careless`, which is freely available and open source (https://github.com/rs-station/careless). We explore the benefits of this approach, specifically in the context of optimizing time-resolved difference peaks, in this work.

The main advantage of variational inference is flexibility. Consequently, software like `careless` should not be thought of as a push-button scaling solution. Rather, it is a modeling toolkit which can be used to extract the most signal possible from a time series. Doing so requires the user to identify the proper set of model hyperparameters to maximize the desired signal. Extensive hyperparameter searches can be computationally prohibitive. Therefore, we offer some lessons from our own experience to help decrease the burden of finding the optimal hyperparameters.

## II. METHODS

### A. Experimental Design

All diffraction data were collected from DJ-1 microcrystals (∼25 *µ*m in size, grown as previously described^5,34^) at APS BioCARS 14-ID-B^12,13^ with a custom sample cell coupled to a microfluidic mixer^35^. Briefly, the DJ-1 microcrystals first pass through a microfluidic mixer where the substrate, methylglyoxal, is rapidly diffused into the crystals for reaction initiation. The freshly mixed crystals continue to flow into the sample cell observation region for data collection. The flow speeds were adjusted to match the 3.6 *µ*s exposure time at a 10 Hz repetition rate so that fresh crystals intersect the X-ray interaction region for every frame. Flow rates and the position of the X-ray beam were adjusted to get different timepoints within the same mixer. Diffraction images were collected at 3, 5, 10, 15, 20, and 30 seconds post mixing, plus an initial state (0-second, without mixing), yielding 7 datasets total. As previously reported^34^, DJ-1 crystallizes in space group *P*3_1_21 which has an indexing ambiguity.

### B. Data Reduction

Data were initially processed using BioCARS’ Python script^13^ for hit finding as well as indexing and integration of images in parallel with Precognition software (Renz Research, Inc.). Next, indexing ambiguities were resolved for each dataset using a custom program (HEX-AMBI, M. Schmidt, personal communication). All details about processing with `careless` can be found on Zenodo (10.5281/zenodo.10481982). Briefly, the *.ii files output from Precognition were converted to .mtz format using to`_mtz.sh`. Then, a custom program, Scramble, was used to correct the consistency of the indexing convention across all datasets (the source code for Scramble is available on GitHub, https://github.com/Hekstra-Lab/scramble/). In this work, we used commit 746597266febb2e8dcf1a0728abe77a47e804bc8 for Scramble and commit 389b3e8084cb5a443a90a3481d80d4197b5b02b3 for `careless.` In addition to being freely available from GitHub, the source code for both of these programs is archived in our Zenodo deposition. Careless^33^ was used to merge and scale the data.

### C. Ablation Study

In order to assess the impact of our `careless` hyperparameter choices on performance, we performed an ablation study whereby we disabled one hyperparameter at a time. The ablations were achieved by making the changes detailed in Figure 1. Dataset quality was assessed with the active site difference peak height, R-factors, *CC*_*hal f*_, *CC*_*work*_*/CC*_*f ree*_, and *CC*_*pred*_. These results are presented in Table I. The active site peak heights were calculated from difference maps between each timepoint post mixing with the initial, 0-second dataset. The average difference peak height and standard deviation across all timepoints for each merge condition is shown in Table I. The R-factors and *CC*_*work*_*/CC*_*f ree*_ are reported for the 0-second timepoint. *CC*_*hal f*_ and *CC*_*pred*_ are reported over all timepoints.

**FIG. 1.**
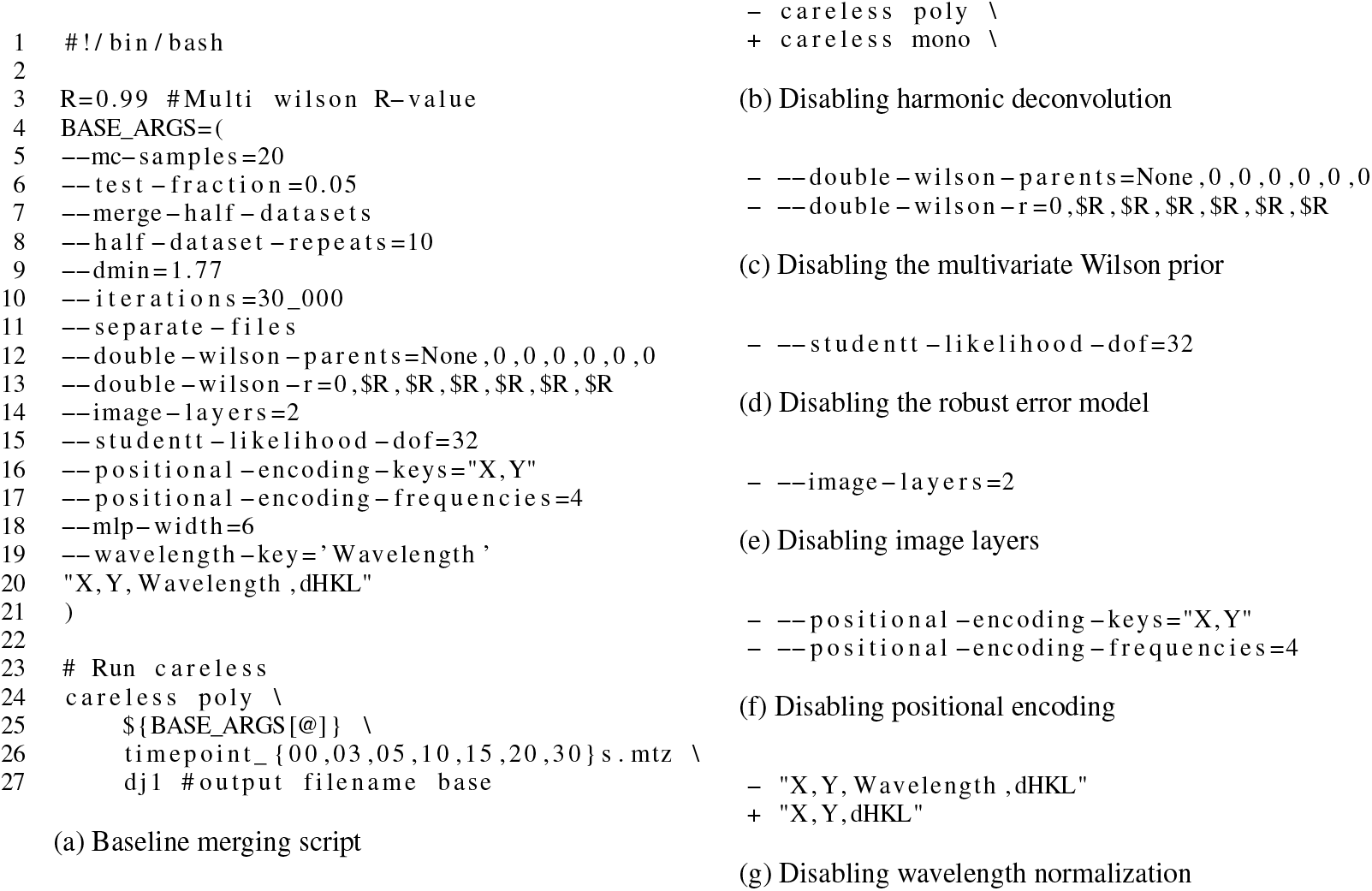
`careless` commands used in this work.

**TABLE I.**
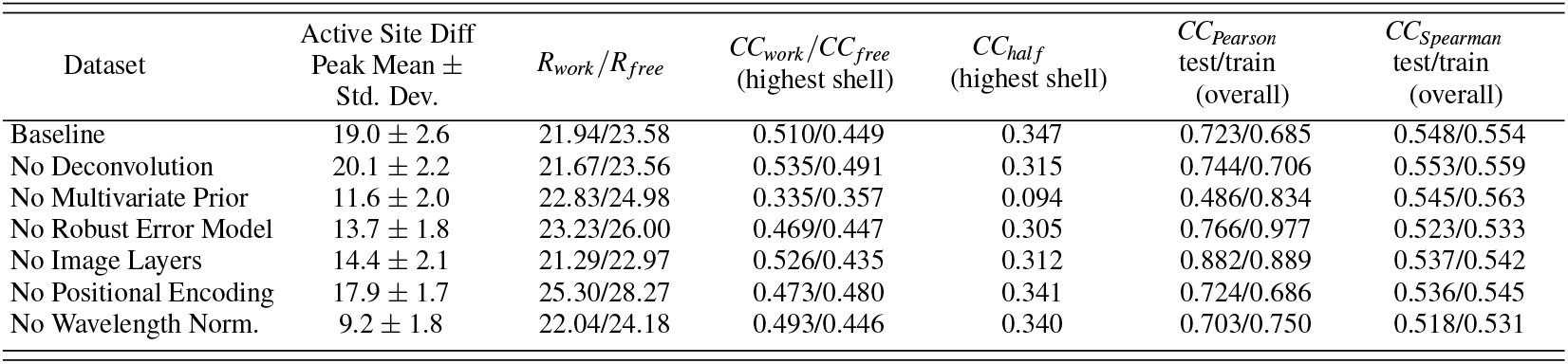
Ablation Studies Results. The average difference peak height and standard deviation across all timepoints for each merge condition is shown. The R-factors and *CC*_*work*_*/CC*_*f ree*_ are reported for the 0-second timepoint. *CC*_*hal f*_ and *CC*_*pred*_ are reported over all timepoints.

### D. Calculation and Quantification of Time-Resolved Difference Maps

We estimated phases for difference map peaks by refinement of a ground state model against the 0-second timepoint data. We restricted refinement to rigid body and individual, isotropic atomic displacement parameters in the PHENIX^36^ software package. We used the `rs.diffmap` function available in the `rs-booster` package for the reciprocalspaceship^37^ Python library. These difference maps use the model phases from PHENIX, and the merging output from careless. We quantified difference map peaks using `rs.find_peaks` which relies on GEMMI^38^. Uncertainties in the peak heights were estimated as the standard deviation from repeated refinement in PHENIX with different random seeds.

## III. CARELESS ABLATION STUDIES

Modern machine-learning algorithms are complex and contain many settings which control their performance. These “hyperparameters” are fixed during model training, and so must be selected or “tuned” by iteratively re-fitting a model with different settings to achieve the best results. With many settings to choose, it is generally impossible to explore the entirety of hyperparameter space for a specific problem. The reality is that determining a successful protocol for training a deep-learning model often relies on a mixture of intuition and one-dimensional hyperparameter searches.

Here we present a protocol for merging time-resolved DJ-1 data based on one-dimensional hyper-parameter searches and our developed intuition. In order to rationalize the impact of each decision involved in selecting our protocol, we conducted an ablation study wherein we disable one model feature at a time relative to the optimized protocol which we will refer to as “baseline”. This method is frequently employed in the deep-learning literature. A famous example of this principle is summarized in Figure 4a of Jumper *et al*. ^39^. The parameters used in our baseline protocol are listed in Figure 1a which uses the same syntax as the `careless` command line interface.

The ‘best’ parameters for merging with `careless` remains an active field of research and may be dataset-specific. Both the ablation study and hyperparameter sweep allowed us to assess the impact of each hyperparameter on the success of merging. We hope these results help future users build an intuition for the most critical parameters, demonstrate a sensible way to approach screening parameters, and share practical advice on how to evaluate the results.

We used the following metrics to judge merged dataset quality: mean and standard deviation of active site difference peak across all time points, *R*_*work*_*/R*_*f ree*_, *CC*_*work*_*/CC*_*f ree*_, *CC*_*hal f*_, and *CC*_*pred*_. Most of these parameters are standard crystallographic figures of merit. *CC*_*pred*_ is used to assess overfitting by evaluating the correlation between predicted and observed reflections^33^, which is an important aspect of hyperparameter selection in any machine-learning model. More specifically for *CC*_*pred*_, we report two approaches, Pearson’s correlation with inverse variance weights (*CC*_*Pearson*_) and Spearman’s rank correlation (*CC*_*Spearman*_). We report these coefficients based on the training data and a set of test reflections which were used to train the the scaling model. Poor performance on the test data is suggestive of overfitting which is an indicator of suboptimal hyperparameter values^40^ (chapter 7). All of our results are summarized in Table I, for details on calculating each metric, see the Methods section. The outcome of each ablation study and, when applicable, one-dimensional parameter sweeps, are presented below in more detail.

### A. Harmonic Deconvolution

One crucial consideration with polychromatic diffraction is that reflection observations need not correspond to exactly one Miller index. In fact, the Laue geometry maps all the reflections from a particular central ray onto the same point on the detector. A central ray is defined as the set of reflections which lie on a ray extending outward from the origin of reciprocal space. For instance, the three reflections indicated in Figure 2 all share the central ray passing through reflection *h* = 0, *k* = 1. These three reflections are called harmonics.^41,42^ In the experiment, they will be stimulated by different wavelengths of X-rays. This can be seen by noting the difference in the magnitude of their scattered beam wavevectors, that is by the length of the arrows in Figure 2. The corresponding wavelengths are tabulated in the table to the right of the figure. Despite the difference in the magnitudes of the scattered wavevectors, the direction is shared. That is, the colored lines in Figure 2 are parallel. In reality, there is only one crystal to serve as the origin for these vectors. Therefore, the reflections end up precisely superposed on the detector. This phenomenon of harmonic overlap is distinct from the spatial overlaps which occur frequently in Laue crystallography^42,43^. Spatial overlaps can be resolved during integration by profile-fitting algorithms. However, harmonic overlaps must be deconvolved during scaling^44^. `Careless`^33^ implements one such algorithm for harmonic deconvolution. In order to assess the impact of harmonic deconvolution on the analysis of our time-resolved data, we ran careless in monochromatic mode using the `careless mono` subprogram in place of the `careless poly` subprogram typically applied to Laue data (Figure 1b). We found that disabling harmonic deconvolution did not markedly affect our results. Specifically, we saw no significant difference in the active site difference peak height or refinement R-values (Table I). Harmonic deconvolution did lead to a slight improvement in *CC*_*hal f*_ in the highest resolution bin. We attribute the lack of a dramatic difference to the high redundancy of these data as well as the experimental design. Specifically, the current beam parameters at BioCARS (^12^) mean that only a very small fraction (1% or less) of observations are typically harmonics. In general, we still recommend using harmonic deconvolution for Laue diffraction. Nevertheless, it will most likely only provide a substantial improvement for low-redundancy data or higher spectral-bandwidth sources.

**FIG. 2.**
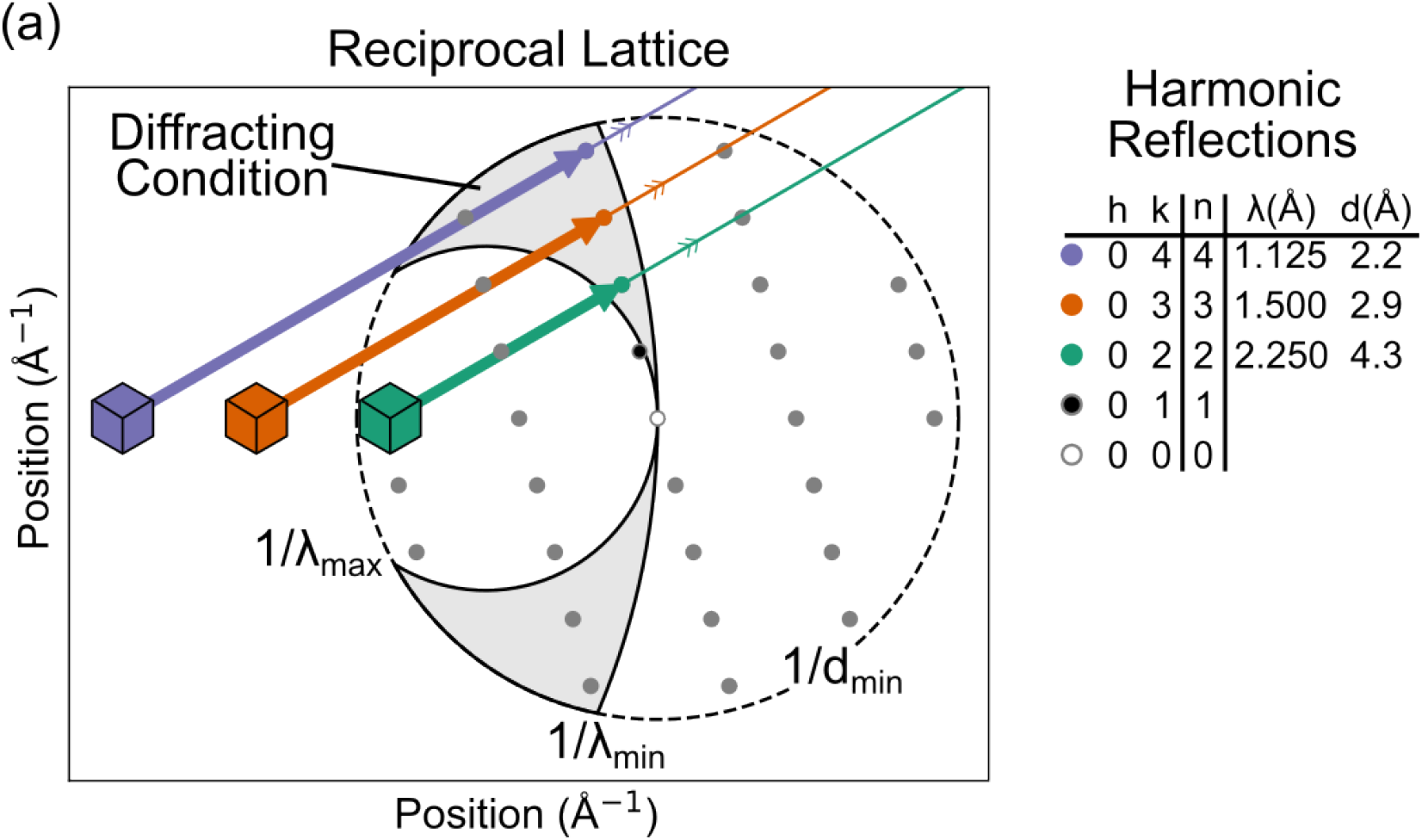
The origin of harmonic overlap in Laue crystallography depicted by the reciprocal lattice for a 2-dimensional crystal. The diffracting condition, shaded grey, is bounded by the three limiting spheres determined by the minimum and maximum wavelength of the spectra and the resolution limit of the sample. Three reflections from the same central ray are on the diffracting condition. Their scattered beam wavevectors, depicted as arrows, have different magnitudes, inverse wavelengths, but are all parallel. The wavelengths and resolutions of these reflections are recorded in the table to the right. Because the scattered beams are parallel, they will arrive at the same location on the detector.

### B. Multivariate Prior

The Bayesian model implemented in `careless` allows the user to express various expectations about the relation between crystallographic data sets through a multivariate prior distribution. To do so, users may specify a conditional dependence structure wherein each data set is dependent on at most one “parent” dataset. Conditional dependence is an appropriate assumption in situations whenever a structure is solved simultaneously with a derivative structure. For each parent-child pair, users specify a value between zero and one for the multivariate prior *r* parameter, which indicates the expected degree of correlation between the two data sets (this distribution is referred to as the “double-wilson” distribution, as it extends the conventional Wilson distribution to the bi- and multivariate case). On the command line, this is done by defining the parent-child relationship with `--double-wilson-parents=None,0,0,0,0,0,0`, where each entry in the list indicates the index of the parent dataset (datasets are indexed in the order in which they are provided to `careless`, starting at 0). In this case, each dataset is a child of the 0^th^ dataset, which corresponds to the 0-second dataset before substrate addition. None indicates that the 0^th^ dataset lacks a parent. Next, the r parameter is set by`--double-wilson-r=0,0.99,0.99, 0.99,0.99,0.99,0.99`, where for each child node the expected correlation with its parent dataset node is set (here 0.990 for all nodes except the 0^th^ (Figure 1c)). Higher numbers indicate a greater expectation of correlation. The multivariate prior *r* is related but not identical to the expected Pearson correlation. As with other hyperparameters discussed here, *r* should be selected on the basis of crossvalidation by assessing the model fit to the training data and a held-out set of test data using the *CC*_*pred*_ measure. The multivariate prior distribution will be described in greater detail in a separate manuscript^45^.

Removing the multivariate prior had a drastic impact on the quality of our analysis. The active site difference peak mean had a 7.4*σ* reduction. Notably, the multivariate prior was the only hyper-parameter to have a considerable effect on *CC*_*hal f*_, thus also impacting the overall resolution of the final dataset as *CC*_*hal f*_ was used as the main criterion to determine the resolution cutoff.

In addition to the ablation study, we also performed a one-dimensional parameter sweep of the prior correlation value (Figure 3). We used the height of the active site difference peak as our main criterion for selecting the optimal value. We found a maximum in difference peak height with a prior correlation value of 0.990 (Figure 3a). At this value, the mean value across all timepoints was 19*σ* whereas the maximum value was 23*σ* (15 s timepoint). Additionally, we far surpass the noise standard of 3*σ* at our optimized value (0.990) and meet or exceed for all of our tested values. The active site difference peak heights, however, were strongly influenced by the prior correlation value (Table I and Figure 3a). We encourage users to carefully screen the prior correlation value for their own datasets to try to find a maximum.

**FIG. 3.**
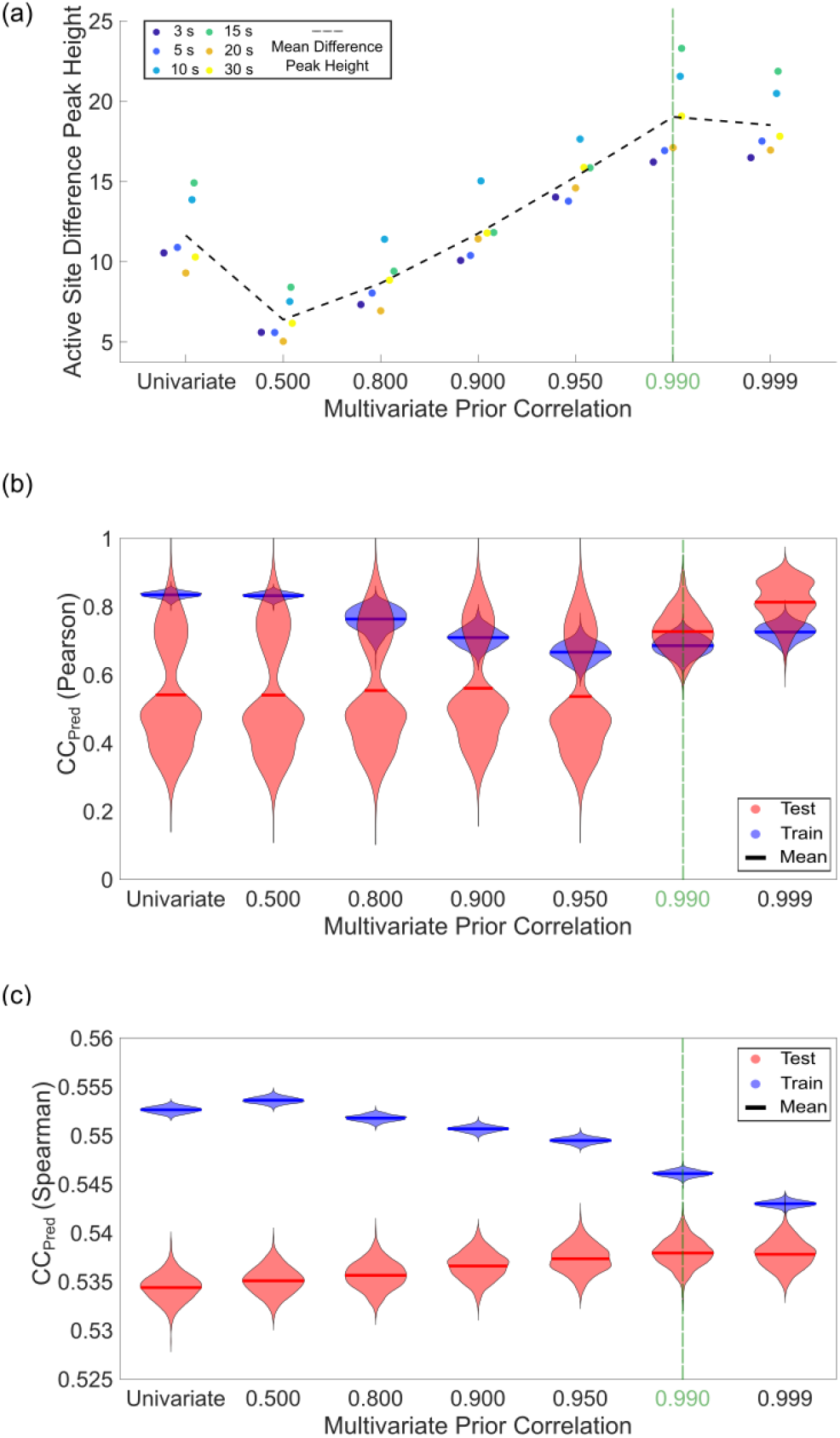
Results of a one-dimensional sweep of the multivariate prior correlation parameter. A. The active site difference peak heights for various multivariate prior correlation values. There is a clear maximum at 0.990. B and C. *CC*_*Pred*_ was calculated to assess overfitting using either the maximum-likelihood weighted Pearson or Spearman correlation coefficients. Careless produces a posterior distribution for each intensity observation. The mean and standard deviation of this distribution are recorded in the `‘*_predictions_#.mtz` files saved after model training. The means of these distributions is typically used to compute *CC*_*Pred*_ or the correlation between observed and predicted intensities. Here, we quantify the uncertainty in *CC*_*Pred*_ using the bootstrap method whereby we resample the predicted intensities recorded in the careless output with replacement 1000 times yielding 1000 estimates of *CC*_*Pred*_ per hyperparameter setting. These bootstrapped estimates are visualized as violin plots. The optimal hyperparameter setting has the highest value while having the smallest gap between test and train. We observe this clearly at 0.990 for *CC*_*Pred*_ (*Pearson*) and at 0.999 for *CC*_*Pred*_ (*Spearman*) indicating that the exact optimum likely lies between the two.

*CC*_*Pearson*_ and *CC*_*Spearman*_ were both computed for crossvalidation. Ideally, these *CC*_*pred*_ indicators should have the highest possible value while having the smallest gap between the test and train set, as this would indicate that the predicted and observed reflections correlate well with each other. In order to estimate the uncertainty of our correlations, we used bootstrapping^46^ to estimate a distribution for both *CC*_*Pearson*_ and *CC*_*Spearman*_. For *CC*_*Pearson*_, we see the smallest gaps between the test and train set median values (Figure 3b) at *r* = 0.990, which corresponds well with our difference peak maximum. Interestingly, *r*-values below 0.990 demonstrate strongly bimodal *CC*_*Pearson*_ distributions which resolve at higher values. At 0.999, the test and train values increase, but the gap between test and train also increases. We interpret this as a less favorable setting. However, we note that the discrepancy in difference peak heights between *r* =0.990 and 0.999 is marginal.

### C. Robust Error Model

A key consideration in fitting models to data is specifying an error model, more specifically how the measurement errors are expected to be distributed. The default choice in careless is that errors follow a normal distribution with a standard deviation determined by the empirical uncertainty estimates from integration. The major drawback of a normally distributed error model is that it is very sensitive to outliers (see Chapter 2.4 of Murphy ^47^). Fortunately, the Bayesian framework used by `careless` is quite flexible and affords some control over the error model. Specifically, `careless` ^33^ provides a robust alternative to the normally distributed error model in the form of a Student’s t-distribution. The Student’s t-distribution has heavier tails than a normal and is therefore more tolerant of outliers^48,49^. The degree of tolerance can be controlled by specifying a parameter of the t-distribution, the degrees of freedom, *ν*. For larger values of degrees of freedom, the model is more sensitive to outliers. As *ν* approaches infinity, the error model becomes equivalent to the normal distribution. In nearly all examples we have encountered, *ν* = 32 has outperformed the normally distributed error model. Therefore, we recommend this setting, which can be implemented on the command line using `--studentt-likelihood-dof=32` (Figure 1d). In our ablation study, we found that the robust, Student’s t-distributed error model was among the strongest determinants of model performance as judged by time-resolved difference peak height (Table I). Specifically, the use of the normal distribution in place of the robust error model resulted in a 5.3*σ* decrease in the mean difference peak height.

We performed a one-dimensional parameter sweep of *ν* to determine if 32 was the optimal value for our data (Figure 4). This was implemented by simply changing `--studentt-likelihood-dof` = to either 4, 8, 16, or 64. We used the active site difference peak heights and both *CC*_*Pearson*_ and *CC*_*Spearman*_ as the main metrics to assess the results (Figure 4). We again used bootstrapping^46^ to estimate a distribution for both *CC*_*Pearson*_ and *CC*_*Spearman*_. By all these criteria it is immediately clear that the Student’s t distribution outperforms the normal distribution, just as demonstrated in Dalton et al., 2022^33^. Upon closer inspection, it is evident that 32 is the best value for the degrees of freedom of this dataset. Although the active site difference peak heights are not strongly affected by *ν*, there is a moderate maximum at 32 (Figure 4a). For *CC*_*Pearson*_, 32 has the smallest difference between the mean test and train and has the train set with the least variability (Figure 4b). For *CC*_*Spearman*_, there is a slight maximum in the test and train mean values and the smallest gap between the test and train sets at 32 (Figure 4c). Overall, this demonstrates that utilizing a few figures of merit is a good practice for selecting values from one-dimensional parameter sweeps. We particularly recommend this combination of using the active site difference peak heights as the main criteria and then both *CC*_*Pearson*_ and *CC*_*Spearman*_ as secondary criteria.

**FIG. 4.**
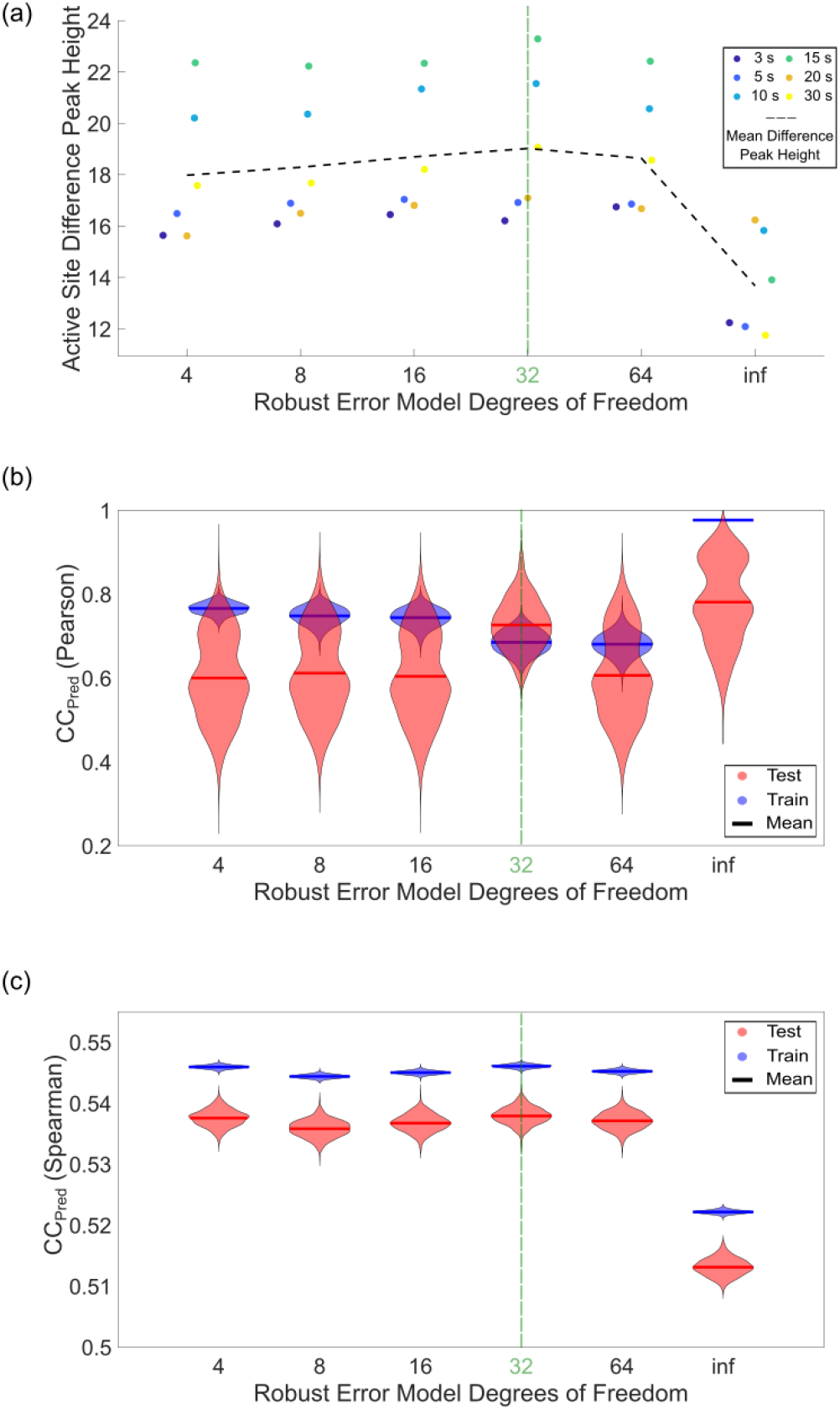
The results of the one-dimensional sweep of the Student’s t degrees of freedom, *ν*, on active site difference peak heights and *CC*_*Pred*_. A. The active site difference peak heights have a slight maximum at 32. B. *CC*_*Pearson*_ has the the smallest gap between test and train at 32, which are the optimal results. C. *CC*_*Spearman*_ is an alternative metric to assess overfitting. It has the best results at 32 with the highest overall value, albeit by a small margin, and the smallest gap between test and train. The distributions visualized in the violin plots were generated by the bootstrap method described in the caption of Figure 3.

### D. Image Layers

By default, `careless` uses a scaling model with purely global parameters. This can be inappropriate for serial crystallography applications wherein each diffraction image typically corresponds to a separate crystal. Variation in the size and quality of samples require different scaling corrections. In such situations, it is important to allocate some of the parameter budget to local parameters which are able to make corrections to each sample independently. In `careless`, this can be done by using image layers. Image layers are neural network layers which have separate parameters for each image in the dataset (see Figure 5a of ref.^33^). The appropriate number of image layers for a dataset can be determined by crossvalidation. In this study we used 2 image layers which we have found to be sufficient in most serial-crystallography cases. This was implemented by using the `--image-layers=2` command line argument (Figure 1e). Crucially, we have never seen 2 image layers lead to overfitting for a serial dataset. In the single-crystal scenario, we have observed cases where image layers can be detrimental to difference map inference. Regarding the DJ-1, serial data, removing the image layers led to a 4.6*σ* decrease in the signal-to-noise of the time-resolved difference peak supporting our assertion that local parameters are beneficial for serial-crystallography data analysis (Table I).

**FIG. 5.**
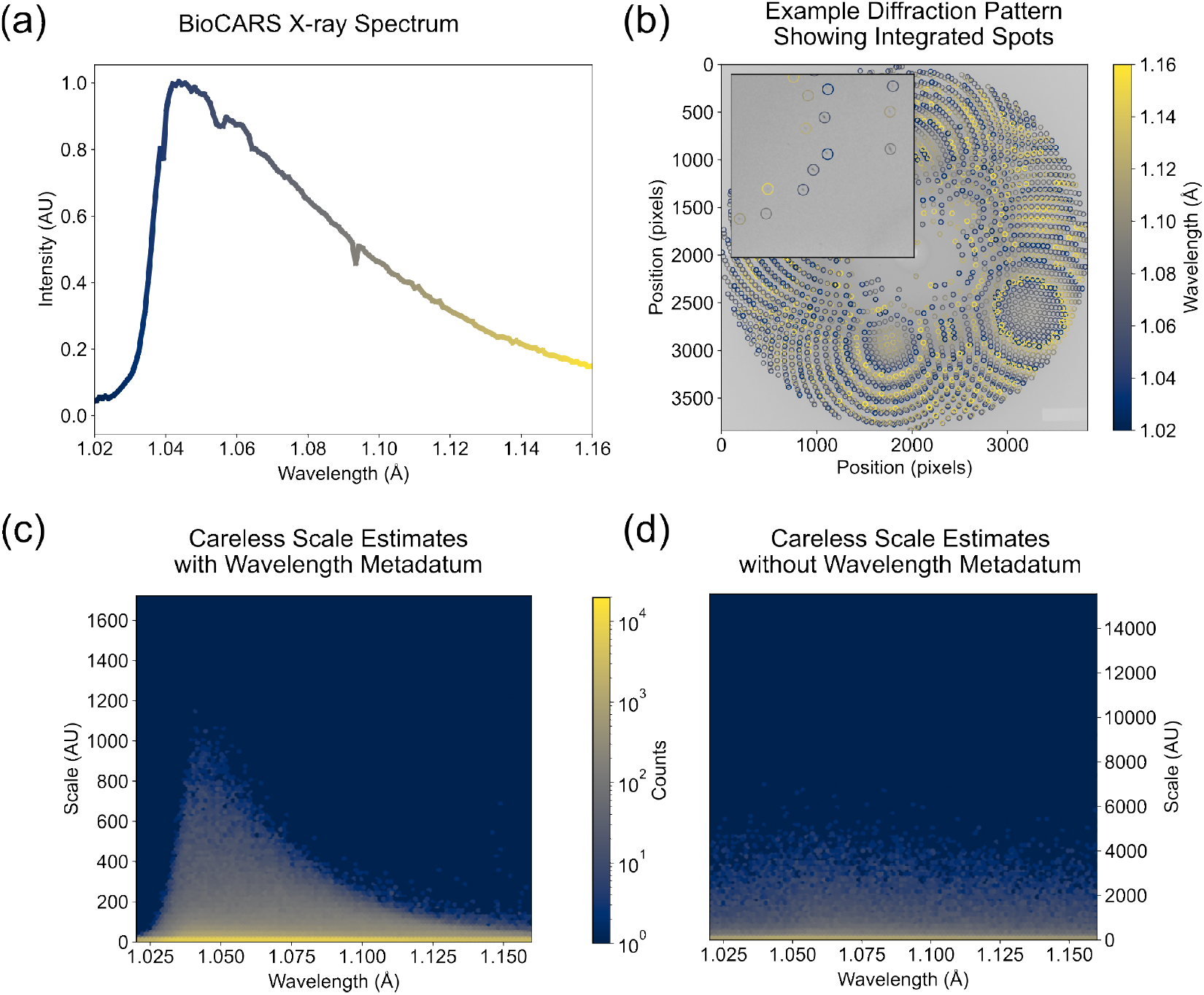
`Careless` infers the spectrum from wavelength metadata. (a) The spectrum of the X-ray beam at BioCARS measured by a channel-cut monochromator. (b) An example diffraction pattern from our DJ-1 data set with the indicated spots circled and colored by the predicted peak wavelength. (c) A two-dimensional histogram of the scale value (systematic error) predicted by `careless` and the peak wavelength predicted by Precognition (Renz Research). The skewed distribution is similar to the BioCARS spectrum indicating that `careless` can infer spectral information. (d) The equivalent two-dimensional histogram produced from the wavelength-normalization ablation study wherein `careless` did not have access to the peak wavelength of each reflection observation.

### E. Positional Encoding

Owing to a variety of effects, the observed intensity of a reflection can vary spatially across the detector^31^. In the case of features such as panel gaps, background scatter, and shadows from the beamstop or other equipment, intensity variations can be abrupt. These sorts of high-frequency variations in can be difficult for neural networks to model^50^. One well-validated solution to this problem is to use a positional encoding strategy^51–53^ whereby coordinates are mapped to a higher-dimensional representation which the scale model can more easily interpret. In `careless`, this is accomplished using the --positional-encoding-keys= command line argument. This parameter takes a set of comma separated metadata keys and instructs `careless` to encode them into a higher-dimensional representation (Figure 1f). The dimensionality of the encoding is controlled by the `--positional-encoding-frequencies=` argument (Figure 1f). In our most successful protocol, we used positional encoding of the detector coordinates of each reflection observation. We found this gave a modest, approximately 2*σ*, increase in difference peak height (Table I). Interestingly, removing positional encoding negatively affected both *R*_*work*_*/R*_*f ree*_ and *CC*_*work*_*/CC*_*f ree*_, so its main benefit seems to contribute to absolute scaling accuracy rather than the accuracy of structure factor differences.

### F. Wavelength Normalization

At synchrotron light sources, it is often necessary to use a polychromatic X-ray beam in order to record time-resolved diffraction at sufficient temporal resolution for biological processes. BioCARS 14-ID beamline at the Advanced Photon Source provides a polychromatic beam with sub-microsecond time-resolution (best time resolution at this beamline is determined by a duration of the single X-ray pulse)^12,13^. Owing to the nature of synchrotron radiation, the BioCARS beam exhibits a characteristically skewed undulator spectrum (Figure 5a). The relatively large bandwidth of the beam provides significantly increased photon flux compared to monochromatic beamlines and consequently much shorter exposure times. It also increases the number of Bragg peaks observed in each image (Figure 5b). However, the spectrum bandwidth introduces an additional challenge in data analysis. In particular, the basal flux differs by wavelength. Therefore, the empirical intensity of each reflection depends on the wavelength at which it is observed on a particular image. In order to model this, a straightforward approach can be used in `careless`. Namely, the empirical wavelength of each reflection observation is computed from the experimental geometry used at integration. Then this wavelength is provided to the `careless` scale model during merging. On the command line, this is done by including `‘Wavelength’` as a metadetum and specifying the key`--wavelength-key=‘Wavelength’` (Figure 1g). Our experiments indicate that `careless` is able to approximate the spectral nature of the beam. This can be best visualized as a histogram of the scale values estimated by careless as a function of wavelength (Figure 5c). As can be seen, the inferred scales recapitulate the characteristic peak and long tail of the pink-beam spectrum. Removing the wavelength metadatum from the `careless` scaling model leads to a featureless dependence of scale on wavelength (Figure 5d) indicating that `careless` is not able to latently infer the wavelength from the remaining metadata. Furthermore, removing the wavelength leads to the worst performing model in our ablation study with time-resolved difference peaks diminished by roughly 10*σ* relative to our baseline.

## IV. DISCUSSION

We demonstrated the importance of various `careless` hyperparameters for getting high-quality merged datasets and especially for capturing time-resolved differences. In particular, we found that wavelength normalization had the largest impact on difference peak heights. This is due to our use of a wide bandwidth 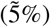 X-ray beam. The next most important parameter, as judged from difference peak heights, is the use of a multivariate prior. For our dataset, we found a prior correlation of *r* =0.990 to be the optimal value, but in general we found the specific value to be quite influential. It is possible that the ideal value may vary for different datasets and types of time-resolved experiment modalities. It is strongly recommended to do a one-dimensional sweep of the multivariate prior to determine which value is best for your specific use case. *CC*_*pred*_-based crossvalidation measures can be unreliable for the multivariate prior. In the limit *r* → 1, the structure factor differences between time points are forced to zero. This equates to learning a single set of structure factors to represent the average across all the time points. Therefore, higher *r*-values can be thought of as reducing the effective parameter count of the model which leads to less apparent overfitting and superior *CC*_*pred*_ values. This indicates a shortcoming of *CC*_*pred*_ in determining the optimal *r*-value. We caution users that for this scenario, *CC*_*pred*_ should not be trusted uncritically. If real-space measures of performance are available, such as the difference peak heights, this is one place where it would be appropriate to rely on those rather than assessments of model fit.

Using a robust error model (the Student’s t likelihood with *ν* = 32) and implementing image layers (with a value of 2) are additional settings which may improve difference map peak heights. These specific values have been consistent for the serial crystallography datasets to which `careless` has been applied thus far^33^ so they are a good starting point. Given the consistency in the performance of these values, one-dimensional sweeps are not necessarily vital for these hyperparameters, especially when using the default parameters for the neural network depth and width (`--mlp-width=10` and --mlp-layers=20). Our `mlp-width` was kept at 6 due to limits on our accelerator card and the large number of reflections across the seven datasets. We recommend you increase `mlp-width` to the maximum value supported by your accelerator card.

Positional encoding had only a modest impact on the difference peak heights, but it had a larger impact on standard crystallographic figures of merit, like *R*_*work*_*/R*_*f ree*_ and *CC*_*work*_*/CC*_*f ree*_. Thus, it is still an important parameter to utilize. Harmonic deconvolution had minimal effects on the analysis quality. This is expected on the basis of the experimental design in this case, in which only a small fraction (<1%) of reflections have contributions from multiple harmonics.

The resolution limit, `dmin`, of the data also needs to be set for every iteration of `careless`. We found that the maximum resolution attainable, based on a *CC*_*hal f*_ threshold of 0.3, was dependent on the hyperparameter values we used. Our baseline dataset has 1.77 Å resolution across all timepoints. We also processed our data with CrystFEL^54,55^. Resolution cutoff for structure refinement for each timepoint in this cases was determined based on >0.2 value of *CC*_*hal f*_ and had a slightly different value for each time point, but the average resolution was 1.98 Å. We attribute this 0.21 Å gain in resolution to the structure imposed by the multivariate prior. This is clearly demonstrated by the degradation of highest-shell *CC*_*hal f*_ exhibited in the multivariate prior ablation (Table I). Since it is ideal to have as high resolution data as possible to interpret time-resolved differences, we see this as a strong argument in favor of using `careless` for time-resolved crystallography data.

Based on our experience, we recommend the following protocol. Starting from an appropriate baseline configuration, such as the one presented in Figure 1a,

1. Determine the resolution cutoff using *CC*_*hal f*_
2. Sweep the multivariate prior correlation (*r*) and determine the optimal value based on realspace performance measures
3. Determine the optimal value of *ν*, the Student’s t likelihood degrees of freedom, based on *CCpred*
4. Re-determine the resolution cutoff using *CC*_*hal f*_

For the first resolution cutoff test (1), we suggest starting with the resolution determined from a standard program, such as CrystFEL^54^, Precognition (Renz Research, Inc.), DIALS^56^, or the Daresbury Laue Suite^57^, and then increase this in small increments determined by your desired precision until you approach a *CC*_*hal f*_ value of 0.3 in the highest resolution shell. For the one-dimensional sweep of the multivariate prior correlation parameter (2), we recommend the following values: 0.500, 0.800, 0.900, 0.950, 0.990, and 0.999. The optimal value should ideally be selected based on a real-space measure, such as the difference peak height.

Once a suitable multivariate Wilson parameter has been established, the Student’s t distribution *ν* sweep can be performed (3). It is most reasonable to test logarithmically-spaced values of *ν*. For instance, *ν* = 4, 8, 16, 32, 64. Lastly, once these hyperparameters have been optimized for your dataset, it may be possible to gain additional resolution. It may be worthwhile to attempt to extend the resolution (4) in small increments (0.01-0.05 Å) until *CC*_*hal f*_ in the highest resolution shell drops below 0.3.

While performing the aforementioned hyperparameter sweeps we suggest calculating all the figures of merit in Table I to assess your results. Sometimes the various figures of merit can give conflicting results for the overall best dataset. For example, sometimes the *R*-factors would decrease (improved result), but the difference peak heights would also decrease (worse result). Whenever they are available, we suggest using difference peak heights as the criterion for selecting the best strategy. We view *CC*_*hal f*_ and *CC*_*pred*_ as the next two most important metrics. *CC*_*hal f*_ acts as a standard crystallographic metric to help evaluate the overall consistency and resolution of the dataset.

*CC*_*pred*_ is useful to assess overfitting and can be estimated using either Pearson’s or Spearman’s rank correlation coefficient. Which estimator is more accurate is likely to depend on the particular data being analyzed. Pearson’s method admits weights derived from the empirical uncertainties of reflection intensities estimated during integration. Spearman’s rank correlation coefficient is generally more robust to outliers. Therefore the quality of the uncertainty estimates and the frequency of outliers in a dataset will impact the relative accuracy of each metric. More study is needed to assess alternative measures of model to data agreement. However, for the time being, we recommend users consider both *CC*_*Pearson*_ and *CC*_*Spearman*_ for hyperparameter optimization. Ultimately, a decision can frequently be made as to which is superior on the basis of difference peak heights.

Overall, we have presented insights on the key parameters when utilizing `careless` to merge time-resolved serial crystallography data. We have also provided practical advice on how to screen such parameters and how to interpret the results. Although our datasets were collected via Laue crystallography at a synchrotron, we expect the described approach to be broadly applicable across serial crystallography modalities.

## ACKNOWLEDGMENTS

KMD holds a Career Award at the Scientific Interface (CASI) from the Burroughs Wellcome Fund. The Linac Coherent Light Source (LCLS) at the SLAC National Accelerator Laboratory is an Office of Science User Facility operated for the U.S. Department of Energy Office of Science by Stanford University. This work was supported by the U.S. Department of Energy, Office of Science, Basic Energy Sciences under Contract No. DEAC02-76SF00515 HKW is supported by a National Science Foundation Graduate Research Fellowship (to HKW, DGE 2140743). This work was funded in part by the National Science Foundation through STC award #1231306 (awarded to LP), the National Institute of General Medical Sciences of the National Institutes of Health under award number DP2-GM141000 (to DRH), R35GM153337 (to MAW), and R35GM122514 (to L.P.), and the National Institute of General Medical Sciences of the National Institutes of Health under grant number P41 GM118217 (R.H. and V.S.) We thank Peter Zwart for guidance on crystallographic twinning.

## DATA AVAILABILITY STATEMENT

The data that support the findings of this study are openly available in Zenodo at 10.5281/zenodo.10481982

## Notes

### Competing Interest Statement

The authors have declared no competing interest.

